# Cryo-EM Structures of Methanogenic Schizorhodopsins Reveal Divergent Strategies for Proton Transport and Thermal Adaptation

**DOI:** 10.1101/2025.10.20.683367

**Authors:** Iota Ebisu, Tatsuki Tanaka, Yuma Kawasaki, Shunya Murakoshi, Keisei Shibata, Yoshitaka Kato, Masae Konno, Wataru Shihoya, Keiichi Inoue, Osamu Nureki

**Affiliations:** Department of Biological Sciences, Graduate School of Science, The University of Tokyo, Bunkyo, Tokyo 113-0033, Japan; The Institute for Solid State Physics, The University of Tokyo, Kashiwa, Chiba 277-8581, Japan; Department of Signal Exploration, The Sakaguchi Laboratory, Keio University School of Medicine, Shinjuku, Tokyo 160-8582, Japan

**Author notes:** Corresponding authors: W. Shihoya, K. Inoue, and O. Nureki.

## Abstract

Schizorhodopsins (SzRs) are light-driven inward proton pumps found in Asgard archaea, contrasting with most microbial rhodopsins that export protons. Some SzRs, including *Methanoculleus taiwanensis* SzR (*Mt*SzR) and *Methanoculleus sp.* SzR (*M*sSzR), exhibit remarkable heat tolerance, suggesting structural adaptations to extreme environments. Here, we report cryo-electron microscopy structures of *Mt*SzR and *M*sSzR at 2.4 and 2.7 Å resolutions, respectively, revealing distinct mechanisms for proton transport and thermostability. Both proteins share a cytoplasm-facing proton acceptor, yet *Mt*SzR employs a hydrophobic gating mechanism, whereas *M*sSzR lacks this barrier and stabilizes its acceptor through a salt-bridge network. Thermostability also diverges: *Mt*SzR relies on proline-rich loops and electrostatic interactions, while *M*sSzR achieves stability through reinforced trimerization and extensive interhelical aromatic packing. Structural and mutational analyses highlight the adaptability of SzRs and provide a framework for engineering robust photoreceptors for optogenetics and synthetic biology.

## Introduction

Rhodopsins are photoreceptor proteins found across bacteria, archaea, and eukaryotes^1–3^. They are broadly classified into three groups: microbial rhodopsins (type 1), animal rhodopsins (type 2), and heliorhodopsins (HeRs), the latter differing substantially in both sequence and membrane orientation from the other two families^4^. Despite their diversity, all rhodopsins share a seven transmembrane helical architecture (TM1–TM7) and contain a retinal chromophore covalently bound to a conserved lysine in TM7 via a Schiff base linkage^5,6^. While the presence of retinal is universal among most rhodopsins, its isomer differs by type: microbial rhodopsins and HeRs typically incorporate all-*trans* retinal, whereas animal rhodopsins use 11-*cis* retinal.

Light absorption triggers isomerization of the retinal chromophore, which drives distinct functional mechanisms in different rhodopsin families. In microbial rhodopsins, all-*trans* to 13-*cis* isomerization initiates a photocycle that supports diverse activities such as ion pumping^7–10^, ion channeling^11–15^, enzyme activity^16^, and phototaxis^17,18^. In contrast, animal rhodopsins act as G protein-coupled receptors, mediating visual signaling via G protein activation ^19^. The ability of microbial rhodopsins to convert light energy into ion gradients has led to their utilization as tools in optogenetics, a technique that allows precise control of neural activity with light^20^.

Unlike microbial and animal rhodopsins, most HeRs lack ion transport activity, and their physiological roles remain unclear^4^. Recent studies, however, have revealed new functions: virally derived V2HeR3 acts as a light-activated proton transporter^21^, and some HeRs interact with enzymes to regulate activity^22,23^. Advances in metagenomic sequencing continue to uncover new rhodopsin families, broadening the scope of light-driven processes in microbial ecology.

In 2020, a new rhodopsin family phylogenetically distinct from previously described groups was identified in the genomes of Asgard archaea—currently classified as Promethearchaeati—microorganisms considered the closest relatives of the last eukaryotic common ancestor ^24–26^. This family was designated schizorhodopsin (SzR) and occupies an intermediate position between microbial rhodopsins and HeRs in sequence-based phylogeny. The crystal structure of SzR4 revealed a topology more similar to microbial rhodopsins than to HeRs, supporting this evolutionary placement^27^.

Functional analyses demonstrated that SzRs act as inward proton pumps, in contrast to the outward proton transporters like most microbial rhodopsins^28–30^. SzR4 also exhibits a shorter intracellular segment in TM6—approximately eight residues shorter than typical microbial rhodopsins—which facilitates proton release into the cytoplasm with minimal structural rearrangement. Based on these structural characteristics and Fourier-transform infrared (FTIR) spectroscopy studies, we previously proposed an untrapped proton release model, in which protons are released directly into the cytoplasm without intermediate trapping by proton acceptors, a mechanism considered specific to SzRs^27,28,30,31^.

Proteins from thermophilic organisms usually exhibit exceptional stability at elevated temperatures. Among rhodopsins, thermophilic rhodopsin (TR) has served as a representative model for investigating mechanism of thermostability in membrane proteins^32^. Subsequently, other outward proton-pumping rhodopsins from thermophilic species were also found to exhibit high thermostability^33,34^. In 2021, SzR genes were identified in high-temperature environments such as mesothermal marine regions and hot springs, suggesting that these SzRs may be adapted for thermal resilience^35^ [14]. We previously compared the thermostability of two such SzRs– one from *Methanoculleus taiwanensis* (*Mt*SzR) and another from a metagenome-assembled genome of *Methanoculleus sp.* (*M*sSzR)– with SzR1, a mesophilic homolog^35,36^. SzR4 shares 38% and 37% sequence identity with *Mt*SzR and *M*sSzR, respectively, while *Mt*SzR and *M*sSzR share 44% identity.

Under denaturation conditions at 85 °C, measured as the lifetime (τ_denature_) of the protein estimated from the decrease in retinal absorption, SzR1 exhibited stability comparable to TR^35^. While *Mt*SzR showed approximately 10-fold greater thermostability than SzR1, *M*sSzR displayed nearly 100-fold higher thermostability, making it the most thermostable rhodopsin reported to date. Notably, whereas all previously characterized thermostable rhodopsins, including TR, function as outward proton pumps, *Mt*SzR and *M*sSzR operate as inward proton pumps^37^. These findings raise fundamental questions regarding the structural determinants of thermostability in SzRs and their relationship to proton transport. Such mechanistic insights are also critical for the rational design of thermostable photoreceptors and enzymes for industrial and synthetic biology applications.

In this study, we determined cryo-electron microscopy structures of *Mt*SzR and *M*sSzR at 2.4 Å and 2.7 Å resolutions, respectively, to investigate the structural basis of thermostability and inward proton transport mechanisms in schizorhodopsins. Comparative structural analysis, combined with mutagenesis, revealed distinct mechanisms that confer extreme thermal stability and enable proton uptake across the membrane.

## Results

### Cyro-EM structures of *Mt*SzR and *M*sSzR reveal conserved architecture and unique features

*Mt*SzR and *M*sSzR were expressed in *Escherichia coli*, purified, and analyzed by single-particle cryo-electron microscopy (cryo-EM). High-resolution structures were obtained at 2.38 Å for *Mt*SzR (reconstituted in nanodiscs) and 2.67 Å for *M*sSzR (in Cymal-4 micelles) (Fig. 1a, b; Supplementary Figs. 1–3). Attempts to reconstitute *M*sSzR into nanodiscs led to fiber-like oligomer formation, which precluded structure determination, necessitating the use of Cymal-4 micelles.

**Fig. 1.**
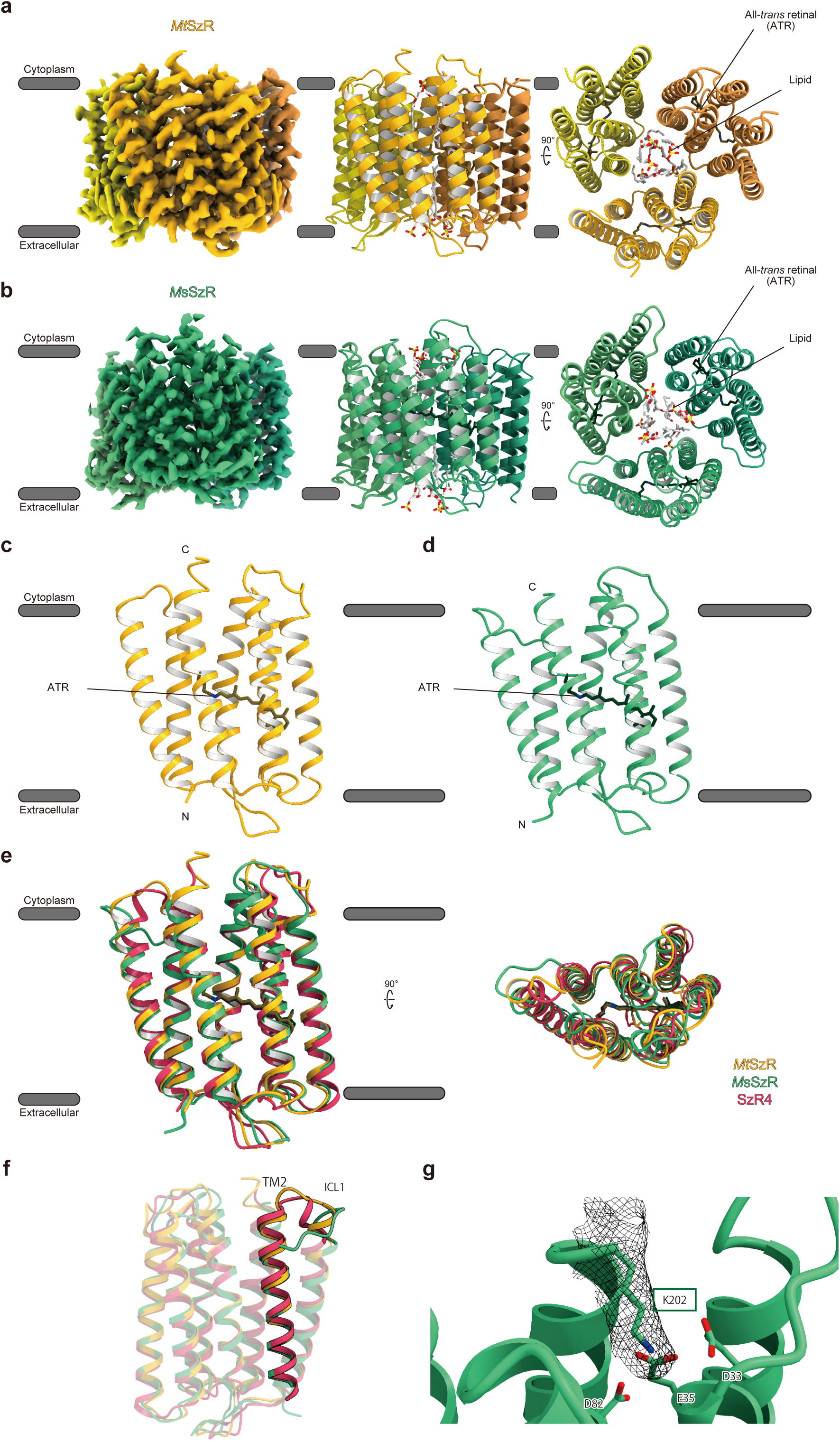
Overall structure of the *Mt*SzR and *M*sSzR. **a, b**. Overall structures of *Methanoculleus taiwanensis* schizorhodopsin (*Mt*SzR) and *Methanoculleus* sp. schizorhodopsin (*M*sSzR), respectively. **c, d**. Monomer structures of MtSzR and MsSzR, respectively. **e**. Superimposition of SzR4 (red), *Mt*SzR (yellow), and *M*sSzR (green). **f**. Comparison of transmembrane helix 2 (TM2) and intracellular loop 1 (ICL1). TM2 of *M*sSzR is one helical turn shorter than SzR4 and *Mt*SzR. **g**. The C-terminal residue K202 of *M*sSzR extends into the space created by the shortened TM2.

Both proteins exhibit the canonical rhodopsin architecture comprising seven transmembrane helices (TM1–TM7) connected by six loops: three extracellular loops (ECL1–ECL3) and three intracellular loops (ICL1–ICL3) (Fig. 1c, d). As in other type 1 rhodopsins and SzRs, the N-terminus faces the extracellular side, and the C-terminus faces the cytoplasm, supported by the previous immunostaining of SzR1^28^, the charged-residue distribution consistent with the positive-inside rule^38^, and the structural superposition with SzR4 (Fig. 1e). ECL1 contains two short antiparallel β-strands, and all-*trans* retinal is covalently linked to a conserved lysine in TM7 via a retinal Schiff base (RSB).

Both *Mt*SzR and *M*sSzR form trimers, stabilized by interactions between TM1 and TM2 of one protomer and TM4’ and TM5’ of an adjacent protomer. Their transmembrane regions align closely with SzR4 (PDB ID: 7E4G), with root-mean-square deviation of 1.17 Å for *Mt*SzR–SzR4 and 1.41 Å for *M*sSzR–SzR4 (Fig. 1e). The greater deviation for *M*sSzR reflects two notable differences: a shortened TM2 and a distinct C-terminal region. TM2 on the cytoplasmic side of *M*sSzR is one helical turn shorter than in SzR4 and *Mt*SzR (Fig. 1f; Supplementary Fig. 4).

Unlike the disordered C-terminal regions in SzR4 and *Mt*SzR, *M*sSzR displays well-defined density, including residue K202, which projects toward the protein core (Fig. 1g). K202 is positioned within the interaction distance of D33, E35, and D82, suggesting potential salt-bridge formation that may contribute to local stabilization. These structural differences—shortened TM2 and an ordered C-terminal region—may underlie the exceptional thermostability of *M*sSzR.

Residues surrounding the retinal chromophore are key determinants of spectral properties and proton transport. In microbial rhodopsins, two negatively charged residues—and only one in SzR4—serve as counterions to stabilize the high pKa of the RSB^27^ [11] (Fig. 2a). Both *Mt*SzR and *M*sSzR contain a single negatively charged residue at the homologous position in SzR4, likely functioning as the counterion (Fig. 2b, c). Aromatic residues Y71 and W154, which are adjacent to the C10–C13 region of retinal in SzR4 and promote all-*trans* to 13-*cis* isomerization, are conserved among 85 SzR homologs, including *M*sSzR and *Mt*SzR, as are the surrounding residues and the hydrogen-bonding network involving water molecules (Supplementary Fig. 5). SzR4 and *Mt*SzR contain three water molecules near the counterion in similar positions, whereas *M*sSzR contains two, one of which is shifted but likely forms an equivalent hydrogen-bonding interaction.

**Fig. 2.**
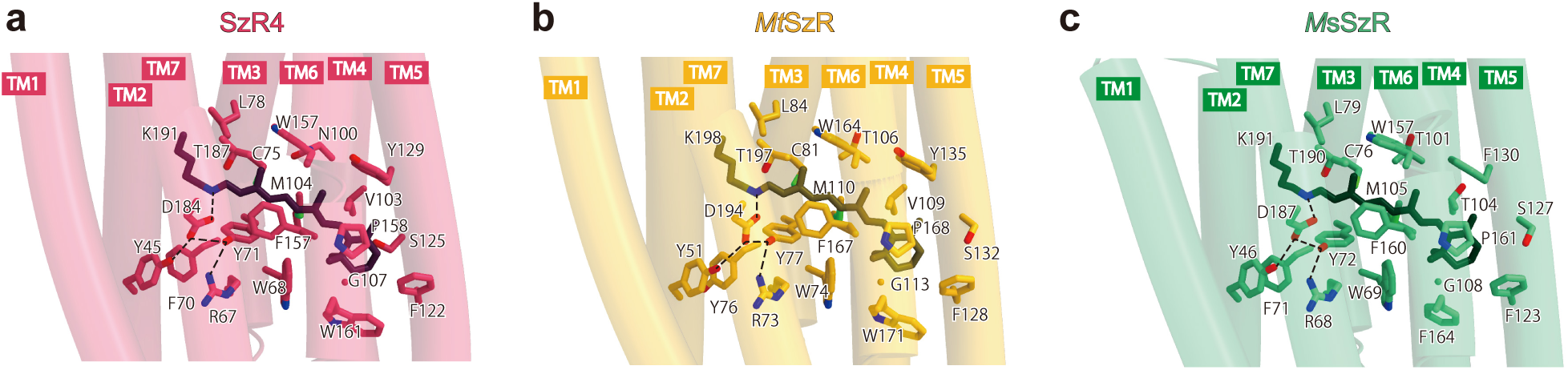
Retinal binding site. Black dashed lines indicate hydrogen bonds. **a-c**. Local environments of the retinal Schiff base (RSB) in SzR4, *Mt*SzR, and *M*sSzR, respectively.

Thus, while the retinal-binding environment—including counterions, aromatic residues, and water-mediated hydrogen-bonding—remains highly conserved among these SzRs (Fig. 2), these features do not account for the enhanced thermostability of *Mt*SzR and *M*sSzR.

### Distinct structural strategies for inward proton transport in *Mt*SzR and *M*sSzR

SzR4 has been proposed to operate via an “untrapped inward proton release” mechanism, in which protons bypass intermediate traps and move directly to the cytoplasm ^27^ (Fig. 3a). In the ground state, the proton acceptor E81 is stabilized by a hydrogen-bond network and shielded from solvent by hydrophobic residues. Upon retinal isomerization from all-*trans* to 13-*cis*, conformational changes in TM3 disrupt these interactions, including hydrogen bonds around E81 (E81–N34, E81–T195) and hydrophobic barriers above (L30, L85) and below (L78) the acceptor. This rearrangement creates a continuous water-mediated pathway between the RSB and cytoplasm, enabling direct proton release driven by the negative charge of the acceptor.

**Fig. 3.**
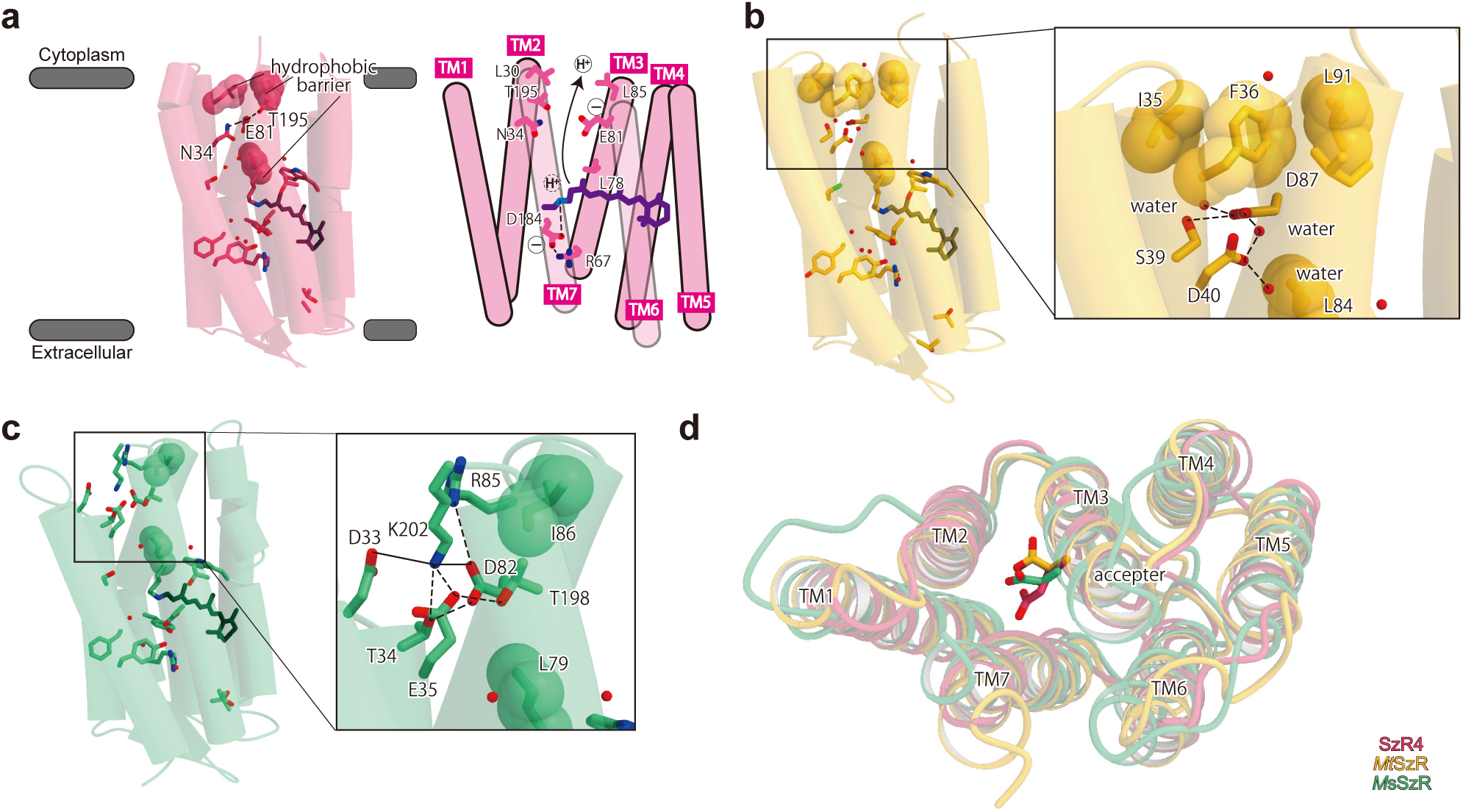
Proton transport. Black dashed lines indicate hydrogen bonds. **a**. Proposed inward proton transport pathway in SzR4. **b-c**. Structures near the proton acceptor in *Mt*SzR and *M*sSzR. **d.** Comparison of acceptor orientations.

*Mt*SzR retains most of the features in this model. Its proton acceptor, D87, is positioned closer to the protein exterior than that of SzR4, and the upper hydrophobic barrier is formed by I35, F36, and L91 (Fig. 3b). Unlike SzR4, where N34 hydrogen bonds with the acceptor, the corresponding D40 in *Mt*SzR is displaced, eliminating the direct interaction with D87. Instead, two water molecules bridge D40 and D87, creating a modified hydrogen-bond network. These differences alter local stabilization of D87 but preserve hydrophobic gating, consistent with an untrapped proton release pathway in *Mt*SzR.

*M*sSzR, by contrast, markedly diverges from this mechanism. Although its proton acceptor D82 also faces outward (Fig. 3c, d), the cytoplasmic hydrophobic barrier is incomplete, formed only by I86, because the cytoplasmic side of TM2, contributing to the barrier in SzR4, is one helical turn shorter in *M*sSzR, which exposes the acceptor to the solvent. Instead of hydrophobic gating, *M*sSzR stabilizes D82 through electrostatic interactions: a salt bridge with C-terminal K202 and an ionic contact with R85 (Fig. 3c). K202 participates in a broader network of salt bridges involving negatively charged residues D33, E35, and D82, suggesting that D33 and E35 may also contribute to inward proton transport. This arrangement likely stabilizes the negatively charged, deprotonated state of D82 and creates an alternative proton-conducting pathway. Given the absence of a continuous hydrophobic barrier, *M*sSzR appears to facilitate inward proton transport via an electrostatic relay rather than the hydrophobic gating mechanism observed in SzR4 and *Mt*SzR. This difference may be related to its characteristic photocycle, which includes direct conversion from the K_2_/L_1_ to the M_3_ without accumulating the L_2_/M_1_ and L_3_/M_2_ states^35,36^. Such mechanistic diversity among SzRs may reflect adaptations to extreme environments, where proton transport must remain efficient under high thermal stress.

### Structural determinants of MtSzR thermostability

To elucidate the structural basis for thermostability of *Mt*SzR, we performed a comparative analysis of its loop regions and mutational studies. In the previous protein engineering, deleting or shortening residues in non-functional loops often enhances stability^39^. However, loop length did not correlate with thermostability among SzRs: SzR1, *M*sSzR, and *Mt*SzR contain 37, 52, and 46 loop residues, respectively. These findings indicate that factors beyond loop length contribute to thermostability.

Proline residues within loops increase protein thermostability by acting as thermal sensors^40^. In TR, prolines are proposed to promote loop integration into hydrophobic environments upon heating, enhancing hydrophobic packing^32^. Additionally, the torsional strain of proline relaxes at high temperatures, facilitating new interhelical hydrogen-bond formation. Notably, *Mt*SzR contains six proline residues in its loops, which is substantially more than SzR4, *M*sSzR, and TR with 1, 1, and 3 prolines, respectively (Fig.4a–d).

To assess the contribution of these prolines to thermostability, we introduced substitutions of prolines observed in *Mt*SzR into SzR1. For the EF loop, where substantial sequence divergence, however, precluded single-site substitution due to the substantial difference in loop length, we generated chimeras replacing the entire loop. SzR1 mutants S62P, D89P, and an EF-loop mimic corresponding to P68, P95, and P151 in *Mt*SzR exhibited increased thermostability compared to wild-type SzR1 (Fig. 4e, g). A combined mutant (S62P/D89P + EF-loop mimic) achieved further thermostability comparable even to wild-type *Mt*SzR, underscoring the additive effects of P68, P95, and EF-loop mimic (Fig. 4f, g).

**Fig. 4.**
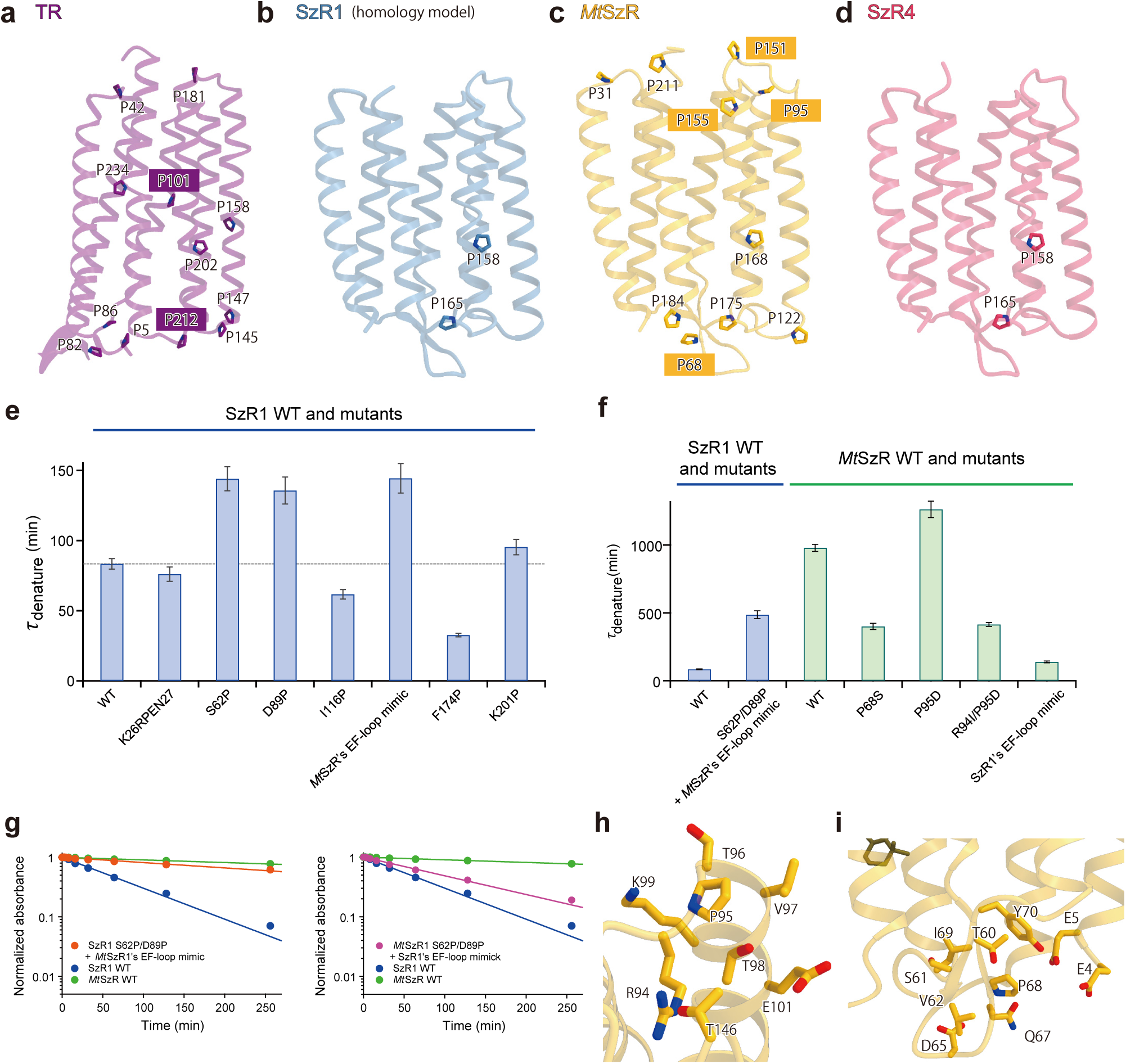
Proline on interhelical loop. **a-d**. Positions of proline residues in SzR1 (homology model based on SzR4), SzR4, *Mt*SzR, and *M*sSzR, respectively. **e**. Thermostability of SzR1 mutants with *Mt*SzR-like proline substitutions. **f**. Thermostability of *Mt*SzR mutants in which prolines were replaced with SzR1-type residues. **g**. Time-dependent changes in retinal absorbance for mutants. **h, i**. Local environments around P68 and P95, respectively. The P95D mutant may form a new salt bridge with nearby R94.

Conversely, reversing proline residues in *Mt*SzR to SzR1-type residues revealed asymmetrical effects. As expected, substitution of P68 with the SzR1-type residue (P68S) reduced thermostability. Unexpectedly, replacing P95 with aspartate (P95D) further enhanced thermostability. This improvement likely arises because the introduced aspartate forms a new salt bridge with the nearby R94 residue, a feature absent in SzR1, thereby enhancing local interactions (Fig. 4h, i). This interpretation was supported by the *Mt*SzR R94I/P95D double mutant, which exhibited reduced stability compared to wild type (Fig. 4f). Overall, these results confirm that loop-localized proline residues, together with local context-specific interactions such as salt-bridge formation, significantly contribute to the exceptional thermostability of *Mt*SzR.

### Structural determinants of *M*sSzR thermostability

To explore the structural basis of *M*sSzR thermostability, we examined its secondary structure and inter-protomer interactions. A notable difference is that TM2 in *M*sSzR is shortened by one helical turn compared to SzR4 and *Mt*SzR, exposing ICL1 toward the cytoplasmic side (Fig. 1g, 5a, b). This exposed loop interacts with TM4 and TM5 of the adjacent protomer, creating a more robust oligomeric interface. At this interface, SzR4 forms three intermolecular hydrogen bonds, *Mt*SzR forms none, and *M*sSzR forms one (Supplementary Fig. 6). These observations indicate that hydrophobic interactions are likely the primary contributors to trimer stabilization, with L30 and F32 in ICL1 playing key roles in contacting the neighboring protomer. The reinforced interface formed by the altered ICL1 may significantly enhance the thermostability of *M*sSzR.

**Fig. 5.**
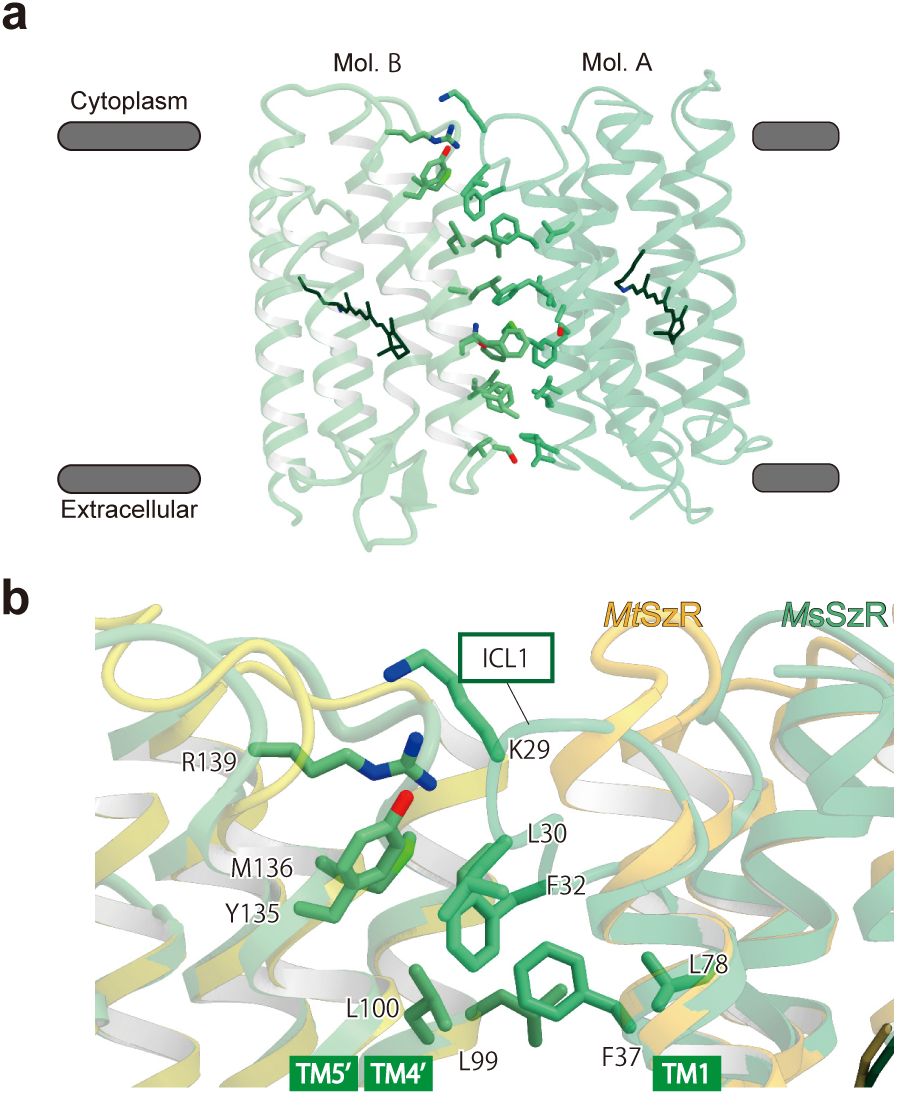
Protomer interface of *M*sSzR. **a**. Side view of the *M*sSzR trimer interface. **b**. Extended ICL1 forms a closer interaction with TM4 and TM5 of the adjacent protomer.

We next considered the potential role of aromatic interactions, which have been linked to protein thermostability in other rhodopsins. In TR, for example, seven aromatic residues contribute to aromatic–aromatic interactions^32^. While the total number of aromatic residues is similar across rhodopsins (40 in TR, 37 in SzR4, 37 in *Mt*SzR, and 34 in *M*sSzR), their distributions differ substantially. While SzR4 shows a higher density of aromatic residues in TM1 and TM7, they are enriched across TM2–TM7 in *Mt*SzR, and *M*sSzR displays a pronounced cluster between TM3 and TM4. This distinct spatial distribution suggests that *M*sSzR may employ a unique stabilization strategy based on localized aromatic packing.

Consistent with this idea, *M*sSzR exhibits the highest number of interhelical aromatic interactions among the SzRs analyzed: 12 in *M*sSzR, compared to 10 in TR, 2 in SzR4, and 7 in *Mt*SzR (Supplementary Fig. 7). This strong correlation between aromatic packing and thermostability implies that such interactions contribute significantly to the thermostability of *M*sSzR. Together, these findings indicate that *M*sSzR achieves thermostability through a combination of reinforced trimerization mediated by an extended ICL1 and an extensive network of interhelical aromatic contacts.

## Discussion

Our *Mt*SzR and *M*sSzR structures provide significant insights into the mechanisms of inward proton transport and thermostability in SzRs, expanding our understanding of functional diversity within this family.

The structural and mechanistic comparisons among SzRs reveal key principles underlying inward proton transport. Both *Mt*SzR and *M*sSzR share a common feature: the proton acceptor is located near the cytoplasmic side (*Mt*SzR: D87, *M*sSzR: D82), and stabilization of its low pKa and negative charge in the ground state is essential for function. In SzR4 and *Mt*SzR, this stabilization is achieved through a hydrogen-bonding network involving nearby residues and water molecules, with surrounding hydrophobic residues shielding the acceptors from solvent. Light-induced structural changes disrupt this network, exposing the acceptor and enabling proton release via an untrapped inward pathway.

In contrast, *M*sSzR employs a distinct strategy. The deprotonated state of D82 is stabilized by a salt bridge with K202 and interactions with R85, rather than by an extensive hydrogen-bonding network. Furthermore, *M*sSzR lacks a complete hydrophobic barrier, leaving D82 partially solvent-exposed even in the ground state. These structural differences suggest that *M*sSzR releases protons through an electrostatic relay rather than the hydrophobic gating observed in SzR4 and *Mt*SzR. Photoactivation likely disrupts the K202-mediated salt-bridge network, allowing protons to move toward the cytoplasm depending on the electrostatic state of the negatively charged acceptor.

Previous mutagenesis studies reported that most SzRs possess a conserved serine near the RSB (∼95%), whereas *M*sSzR has an alanine at this position (A75)^35^. Substitution of this serine in SzR4 (Ser74Ala) abolishes inward proton-pumping activity, suggesting its role in forming the hydrogen-bonding network near the RSB. Although we considered the possibility that water recruitment could compensate for the missing serine in *M*sSzR, our structural model did not identify additional water molecules in this region.

Beyond their proton transport properties, *Mt*SzR and *M*sSzR exhibit remarkable thermostability achieved through distinct structural strategies. *Mt*SzR primarily relies on loop-localized proline residues and context-specific electrostatic interactions, whereas *M*sSzR stabilizes its structure through reinforced oligomerization and extensive interhelical aromatic packing. These contrasting mechanisms underscore the adaptability of SzRs to high-temperature environments and provide a framework for exploring the structural principles underlying their thermal resilience.

In *Mt*SzR, enhanced thermostability is strongly linked to loop-localized proline residues and their functional roles. Mutational analyses demonstrated that P68 and P95 are critical determinants of thermostability. The unexpected P95D salt bridge was found to enhance thermostability, and the disruption of this interaction by the double mutant led to a decrease in thermostability, indicating that the structural integrity at this position is critical for the thermal stability of *Mt*SzR.

*M*sSzR achieves thermostability through a distinct structural strategy involving trimer interface reinforcement and extensive aromatic interactions. The shortening of TM2 exposes ICL1 toward the cytoplasmic side, enabling additional contacts with adjacent protomers and creating a more stable oligomeric interface. Notably, *M*sSzR exhibits the highest number of interhelical aromatic interactions among SzRs. These combined features underpin the exceptional thermal stability of *M*sSzR.

Together, these findings demonstrate that *Mt*SzR and *M*sSzR employ fundamentally different mechanisms to achieve both inward proton transport and thermostability. It suggests that *M*sSzR has followed a distinct evolutionary path from other SzRs. In addition to its remarkable thermal stability, *M*sSzR is known to exhibit greater resistance to acidic conditions^35^. It raises the possibility that multiple selective pressures—including low pH—have influenced its adaptation. By employing a finely tuned salt bridge network to electrostatically stabilize the proton acceptor, *M*sSzR has evolved a compact structure that is resistant to both heat and acidic pH. This diversity highlights the structural plasticity of SzRs and provides a framework for rational engineering of thermostable photoreceptors for optogenetic, industrial, and synthetic biology applications.

## Methods

### Expression and Purification

The genes encoding *Methanoculleus taiwanensis* SzR (*Mt*SzR) and *Methanoculleus* sp. SzR (*M*sSzR) (GenBank ID: DSDF01000045.1), both codon-optimized for *Escherichia coli* expression, were synthesized by GenScript and subcloned into the pET21a(+) vector with an N-terminal His tag. Mutants were generated using the infusion method by designing primers containing the desired mutations with overlapping regions, followed by PCR amplification to obtain plasmids carrying the intended mutations.

The *Mt*SzR and *M*sSzR plasmids were transformed into *E. coli* C41 (DE3) cells. Transformants were selected on LB agar plates supplemented with 100 µg/mL ampicillin, and individual colonies harboring the *Mt*SzR or *M*sSzR expression vectors were isolated. Colonies were picked, and *E. coli* derived from them were pre-cultured in 20 mL of liquid LB medium containing 100 µg/mL ampicillin. The preculture was added to 1 L of TB medium containing 100 µg/mL ampicillin and 8 mL/L glycerol, and cultured at 37°C and 140 rpm until the optical density at 600 nm reached approximately 0.6. Protein expression was induced by the addition of 0.1 mL of 1 M IPTG and 0.1 mL of 0.1 M all-*trans* retinal (ATR), followed by incubation at 37 °C for 4 h. Cells were harvested by centrifugation at 5,000 × g for 10 min.

The harvested *E. coli* cells were resuspended in sonication buffer (20 mM Tris-HCl pH 8.0, 200 mM NaCl, 10% glycerol) and disrupted by sonication. The resulting lysate was subjected to ultracentrifugation at 40,000 rpm for 1 h to collect the membrane fraction as a pellet.

The membrane pellet was resuspended in membrane resuspension buffer (20 mM Tris-HCl pH 8.0, 200 mM NaCl, 10% glycerol) and homogenized using a homogenizer. Solubilization was performed by adding *n*-dodecyl-β-D-maltoside (DDM) to a final concentration of 1% and gently mixing at 4 °C for 1 h. The solubilized mixture was then ultracentrifuged at 35,000 rpm for 20 min, and the supernatant containing the solubilized fraction was collected, while the pellet containing insoluble material was discarded.

Ni-NTA resin was equilibrated using ultrapure water followed by wash buffer (*Mt*SzR: 20 mM Tris-HCl pH 8.0, 500 mM NaCl, 20 mM imidazole, 0.02% DDM; *M*sSzR: 20 mM Tris-HCl pH 8.0, 500 mM NaCl, 20 mM imidazole, 2% Cymal-4). The solubilized supernatant was mixed with equilibrated Ni-NTA resin and gently rotated at 4 °C for 1 h to bind the His-tagged target proteins. The resin was then washed with about 10 column volumes of the respective wash buffer to remove non-specifically bound proteins. The bound target proteins were eluted using elution buffer (*Mt*SzR: 20 mM Tris-HCl pH 8.0, 500 mM NaCl, 300 mM imidazole, 0.03% DDM; *M*sSzR: 20 mM Tris-HCl pH 8.0, 500 mM NaCl, 300 mM imidazole, 1% Cymal-4), and the eluted fractions were concentrated. The samples were loaded onto a Superdex 200 10/300 Increase size-exclusion chromatography column equilibrated with gel filtration buffer (*Mt*SzR: 20 mM Tris-HCl pH 8.0, 150 mM NaCl, 0.03% DDM; *M*sSzR: 20 mM Tris-HCl pH 8.0, 150 mM NaCl, 0.03% DDM, 0.8% Cymal-4) and purified by gel filtration chromatography (Supplementary Fig. 1). For *M*sSzR, the peak fractions were further concentrated using an Amicon Ultra-0.5 centrifugal filter (30,000 MWCO) until the protein concentration reached 3.6 mg/mL, as determined by UV absorbance using a Nanodrop spectrophotometer (Thermo Scientific). The final sample was ultracentrifuged and used for grid preparation.

The peak fractions of *Mt*SzR were mixed with soybean-derived phosphatidylcholine (SoyPC) dissolved in Nanodisc buffer (20 mM Tris-HCl pH 8.0, 500 mM NaCl, 0.5% sodium cholate) and the membrane scaffold protein MSP1E3D1 at a molar ratio of *Mt*SzR:SoyPC:MSP1E3D1 = 4:200:1. The mixture was incubated at 4 °C for 1 h in the dark. Following incubation, 20 mg of Bio-Beads SM2 (Bio-Rad), pre-equilibrated with buffer (20 mM Tris-HCl pH 8.0, 500 mM NaCl), were added and gently rotated at 4 °C. The Bio-Beads were replaced twice at 2 h intervals with equilibrated beads, followed by the addition of 60 mg Bio-Beads and further incubation overnight to remove detergents and reconstitute *Mt*SzR into Nanodisc. After reconstitution, only the supernatant was collected using a narrowed pipette tip, and the sample was ultracentrifuged at 40,000 rpm for 20 min to remove any aggregated proteins. The resulting supernatant was subjected to size-exclusion chromatography using a Superdex 200 10/300 Increase column (GE Healthcare) equilibrated with gel filtration buffer 2 (20 mM Tris-HCl pH 8.0, 150 mM NaCl) (Supplementary Fig. 1). For the *Mt*SzR Nanodisc peak fractions, the sample was concentrated using an Amicon Ultra-0.5 centrifugal filter (30,000 MWCO) until A280 reached 12, as determined by UV absorbance with a Nanodrop spectrophotometer (Thermo Scientific). The final sample was ultracentrifuged and supplemented with digitonin at a final concentration of 0.01% prior to grid preparation.

### Cryo-EM single-particle analysis

3 µL of *Mt*SzR and *M*sSzR Nanodisc samples were applied to glow-discharged holey carbon grids (Quantifoil Au 300 mesh R1.2/1.3) and flash-frozen in liquid ethane using a Vitrobot Mark IV (Thermo Fisher Scientific). Cryo-electron microscopy (cryo-EM) images were collected using a Titan Krios electron microscope operating at 300 kV, equipped with a Gatan K3 Summit direct electron detector. Data acquisition was performed using the EPU software (Thermo Fisher Scientific). Images were recorded at a nominal magnification of 105,000× with a defocus range of −0.6 to −1.6 µm. Each micrograph was captured over a total exposure time of approximately 2 s, at a dose rate of ∼15 electrons per pixel per second, and fractionated into 48 frames. In total, 12,309 movies were recorded for *Mt*SzR and 9,891 movies for *M*sSzR. All movies were acquired in super-resolution mode, binned 2×, dose-fractionated, and subjected to beam-induced motion correction using the implementation in cryoSPARC^41^. The contrast transfer function (CTF) parameters were estimated using the Patch CTF Estimation module in cryoSPARC^42^. Particles were initially picked from a small fraction with Gaussian blob picking and subjected to 2D classification. 8,692,253 particles (*Mt*SzR) and 7,273,694 particles (*M*sSzR) were picked and extracted, followed by 2D classification to remove non convergent particles. A total of 2,128,380 particles (*Mt*SzR) and 394,476 particles (*M*sSzR) were re-extracted with the pixel size of 1.16 Å and curated by heterogeneous refinement^43^.

For *Mt*SzR, 187,792 particles were reconstructed using non-uniform(NU) refinement with C3 Symmetry by cryoSPARC, resulting in a 2.73 Å overall resolution reconstruction of the trimer of *Mt*SzR, with the gold standard Fourier Shell Correlation (FSC = 0.143) In addition, particles were polished on Relion3.1 and selected by 2D classification and NU refinement and then reconstructed using NU refinement with C3 Symmetry and sharpened by cryoSPARC, resulting in a 2.38 Å overall resolution reconstruction of the trimer of *Mt*SzR. The data processing workflow is illusitrated in Supplementary Fig. 2.

For *M*sSzR, the 206,620 particles were reconstructed using NU refinement with C3 Symmetry by cryoSPARC, resulting in a 2.94 Å overall resolution reconstruction of the trimer of *M*sSzR, In addition, particles were polished on RELION 3.1^44^ and selected by 2D classification and NU refinement and then reconstructed using NU refinement with C3 Symmetry by cryoSPARC, resulting in a 2.67 Å overall resolution reconstruction of the trimer of *M*sSzR. The data processing workflows are illustrated in Supplementary Fig. 3.

### Model building and refinement

Initial models of *Mt*SzR and *M*sSzR were built using the crystal structure of SzR4 (PDB ID: 7E4G) as a template. These initial models were roughly fitted into the cryo-EM maps using the modeling program COOT, followed by manual model building to better match the density^45^. The resulting models were refined in real space using PHENIX with secondary structure restraints applied^46,47^. All structural figures were prepared using CueMol (http://www.cuemol.org/ja/) and UCSF ChimeraX^48^. The Statistical values were calculated using MolProbity^49^.

The final model of *Mt*SzR contained residues 4–213, All-*trans* retinal, 13 1,2-diacyl-sn-glycero-3-phosphocholine molecules, and 19 water molecules. The final model of *M*sSzR contained residues 1–202, All-*trans* retinal, 2 1,2-diacyl-sn-glycero-3-phosphocholine molecules, and 23 water molecules.

### Homology Modeling of SzR1

The SzR1 homology model was built based on the crystal structure of SzR4 using SWISS-MODEL^50^.

### Thermostability assay

For thermostability assay experiments, protein expression was carried out with the same method as previously described^35^. After expression, harvested cells were disrupted by the ultrasonic homogenizer (Ultrasonic Homogenizer VP-300 N, TAITEC) in buffer containing 50 mM Tris-HCl (pH 8.0) and 5 mM MgCl_2_. The disrupted cells were collected by the ultra by ultracentrifugation (CP80NX, Eppendorf Himac Technologies) at 125,000 × g for 1 h. The collected cells were homogenized with the potter-type homogenizer and solubilized in buffer 50 mM MES-NaOH (pH 6.5), 300 mM NaCl, 5 mM imidazole, and 1% DDM (ULTROL® Grade; Merck KGaA) for 1h. Ultracentrifugation was at 125,000 × g for 1 h carried out to separate solubilized proteins from insoluble proteins. Solubilized proteins were purified by the affinity chromatography using a Co-NTA affinity column. The resin was washed with buffer containing 50 mM MES-NaOH (pH 6.5), 300 mM NaCl, 27.5 mM imidazole, and 0.1% DDM. The proteins were eluted in buffer containing 50 mM MES-NaOH (pH 6.0), 300 mM NaCl, 500 mM imidazole, and 0.1% DDM. The eluted proteins were dialyzed using buffer 20 mM Tris-HCl (pH 8.0), 150 mM NaCl, 0.02% DDM to remove imidazole.

The thermal stability of both wild-type and mutant forms of *Mt*SzR and *M*sSzR was estimated based on the decrease in visible absorbance. Measurements were performed in a buffer containing 20 mM Tris-HCl (pH 8.0), 150 mM NaCl, and 0.02% DDM. The optical density at the absorption maximum wavelength (λ_max_) of each protein was adjusted to 0.25, and 0.6 mL of the sample solution was incubated at the target temperature (95°C) using a block heater (DTU-Mini, TAITEC). After incubation, the solution was rapidly cooled on ice and centrifuged at 10,000 × g for 3 minutes to remove aggregated protein. The absorption spectra of the supernatant were recorded using a UV-visible spectrophotometer (V-750, JASCO), and the loss of retinal absorbance was estimated from the decrease in absorbance at (λ_max_).

### Data Availability

Cryo-EM Density maps and structure coordinates have been deposited in the Electron Microscopy Data Bank (EMDB) and the Protein Data Bank (PDB), with accession codes EMD-XXXXX and PDB XXXX for *Mt*SzR, EMD-XXXXX and PDB XXXX for *M*sSzR.

## Acknowledgements

We thank K. Ogomori and C. Harada for their technical assistance. This work was supported by JSPS KAKENHI grants 24K23232 (T.T.), 24KJ0909 (S.M.), 24K23073 (Y.K.), 23K05007 (M.K.), 22K19371, 23K24014, 25K02397 (W.S.), 24H02268, 25H00424 (K.I.), 21H05037 (O.N.); AMED under grant number JP233fa627001 (O.N.); the Platform Project for Supporting Drug Discovery and Life Science Research (Basis for Supporting Innovative Drug Discovery and Life Science Research [BINDS]) from AMED under grant numbers JP24ama121002 and JP24ama121012; JST CREST under grant number JPMJCR22N2 (K.I.); Brain Science Foundation (W.S.); MEXT Promotion of Development of a Joint Usage/ Research System Project: Coalition of Universities for Research Excellence Program (CURE) under grant number JPMXP1323015482 (K.I.).

## Author contributions

I.E. performed all of the experiments. S.M assisted with the expression and the purification. grid preparation. T.T assisted with the cryo-EM data collection and single particle analysis. Y.K., K.S and M.K performed thermostability assay and transport assay. The manuscript was mainly prepared by I.E. T.T., and W.S., with assistance from O.N and K.I.

## Competing interests

All authors declare no competing interests.

## Notes

### Competing Interest Statement

The authors have declared no competing interest.

