## Supplementary information for "Cryo-EM Structures of Methanogenic Schizorhodopsins Reveal Divergent Strategies for Proton Transport and Thermal Adaptation"

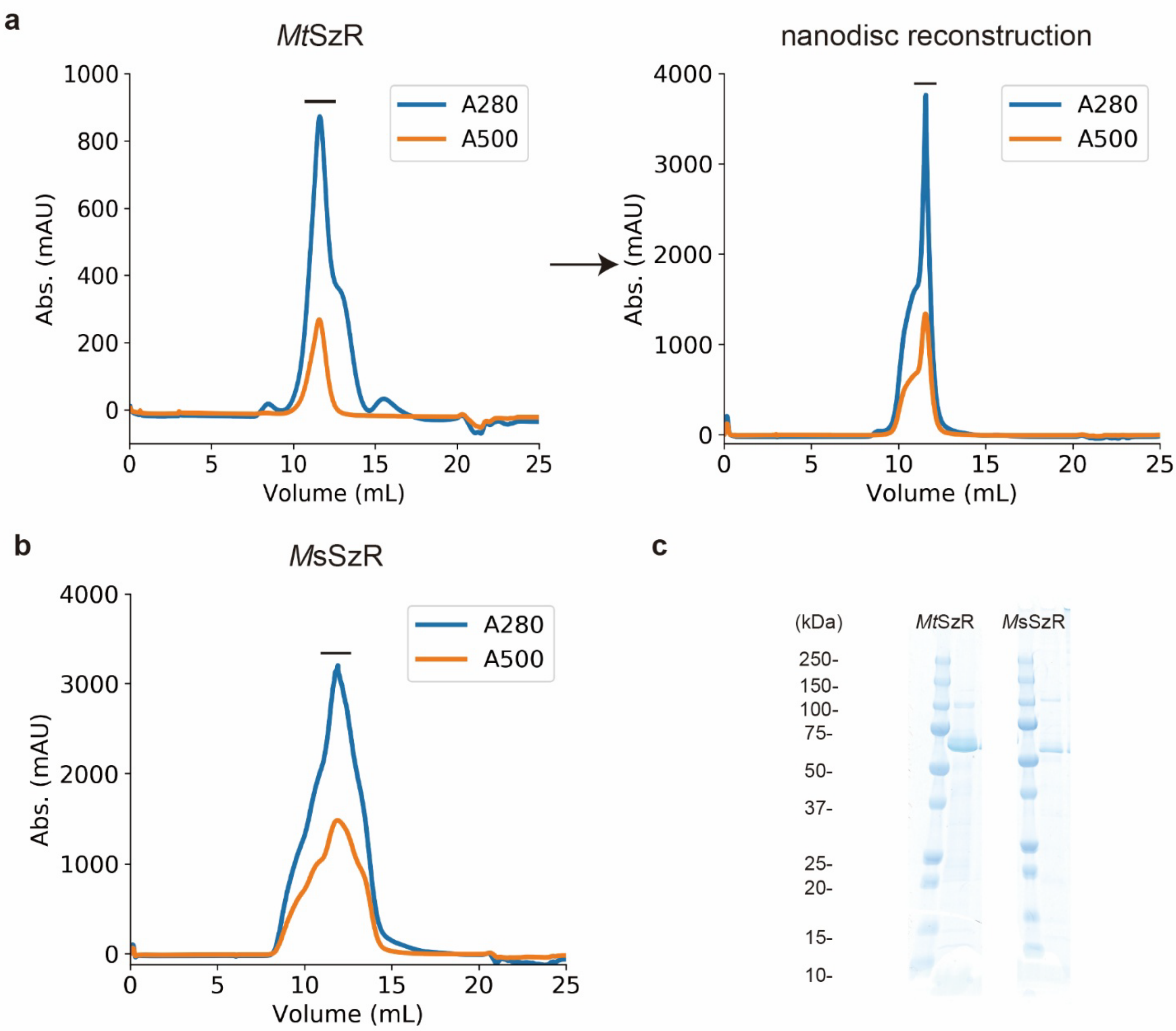


**Figure S1. a**. Chromatogram of *Mt*SzR. The peak fractions indicated by the lines were collected. **b**. Chromatogram of *Ms*SzR. The peak fractions indicated by the line were collected. **c**. SDS -PAGE results of the concentrated samples.

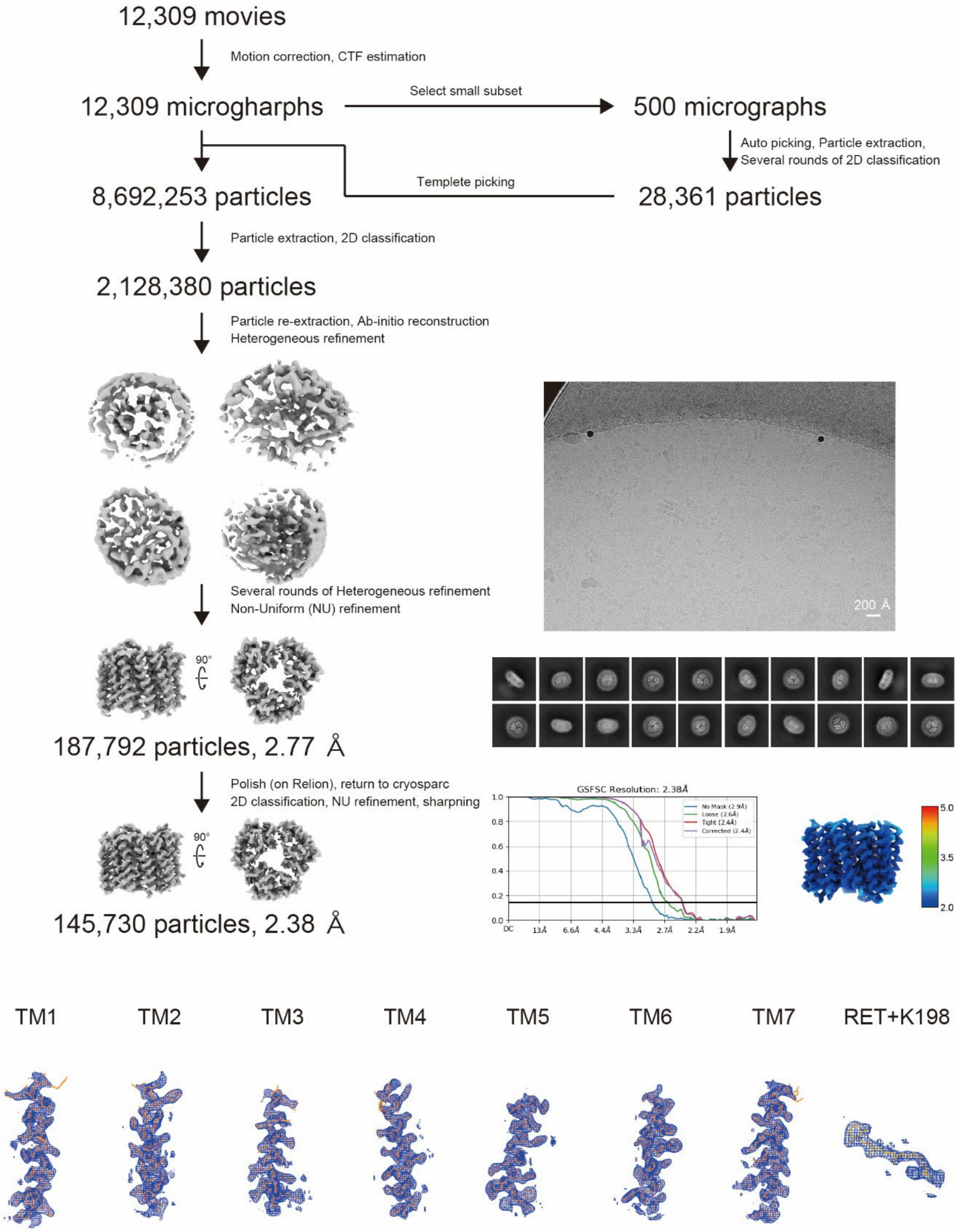


**Figure S2.** Flow chart of the cryo-EM data processing for the *Mt*SzR, representative micrograph, local resolution maps, FSC-curves, and density maps with the atomic model. The density maps were obtained from the NU refinement of the C3 symmetry. Details are provided in the method section.

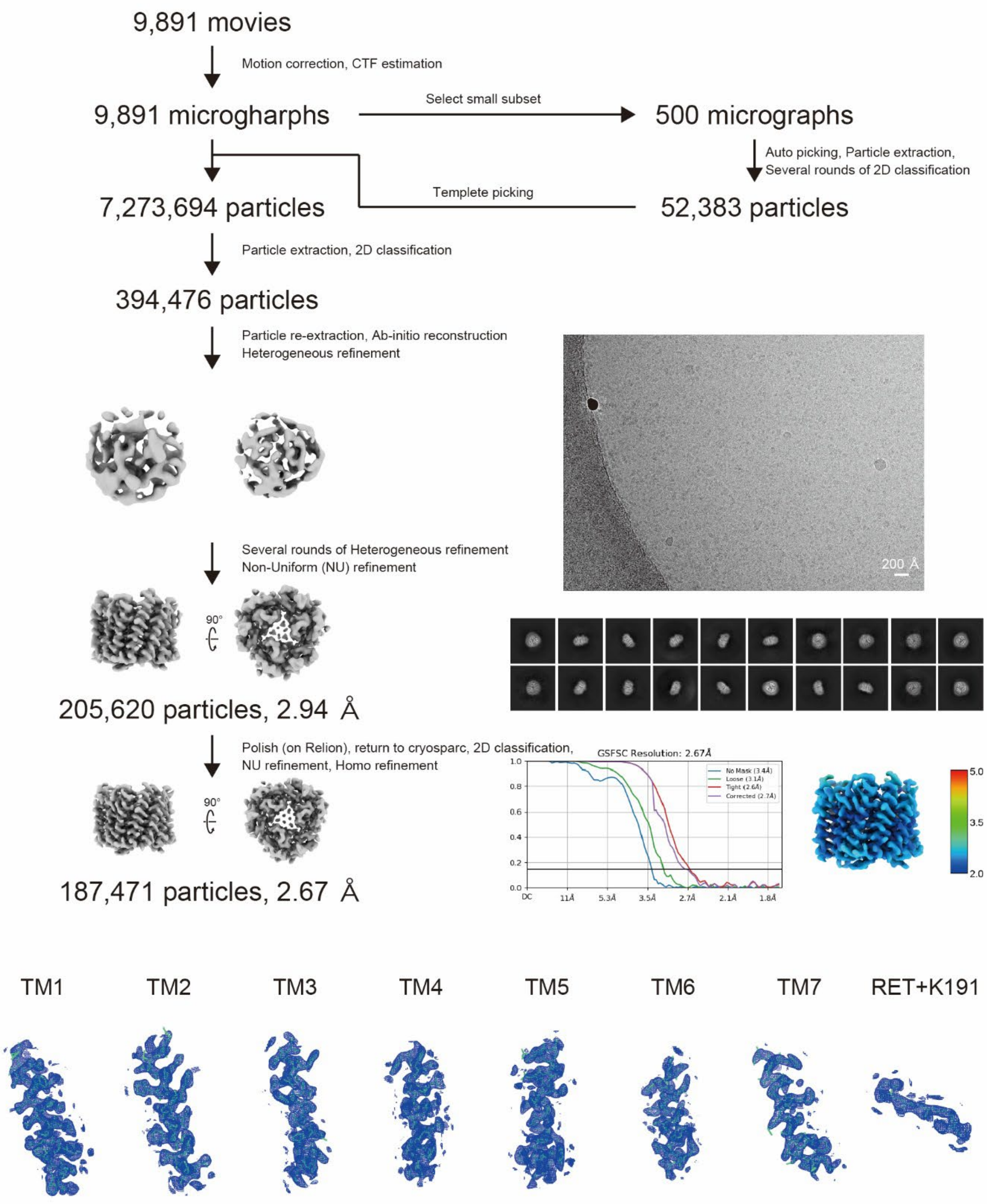


**Figure S3.** Flow chart of the cryo-EM data processing for the *Ms*SzR, representative micrograph, local resolution maps, FSC-curves, and density maps with the atomic model. The density maps were obtained from the NU refinement of the C3 symmetry. Details are provided in the method section.

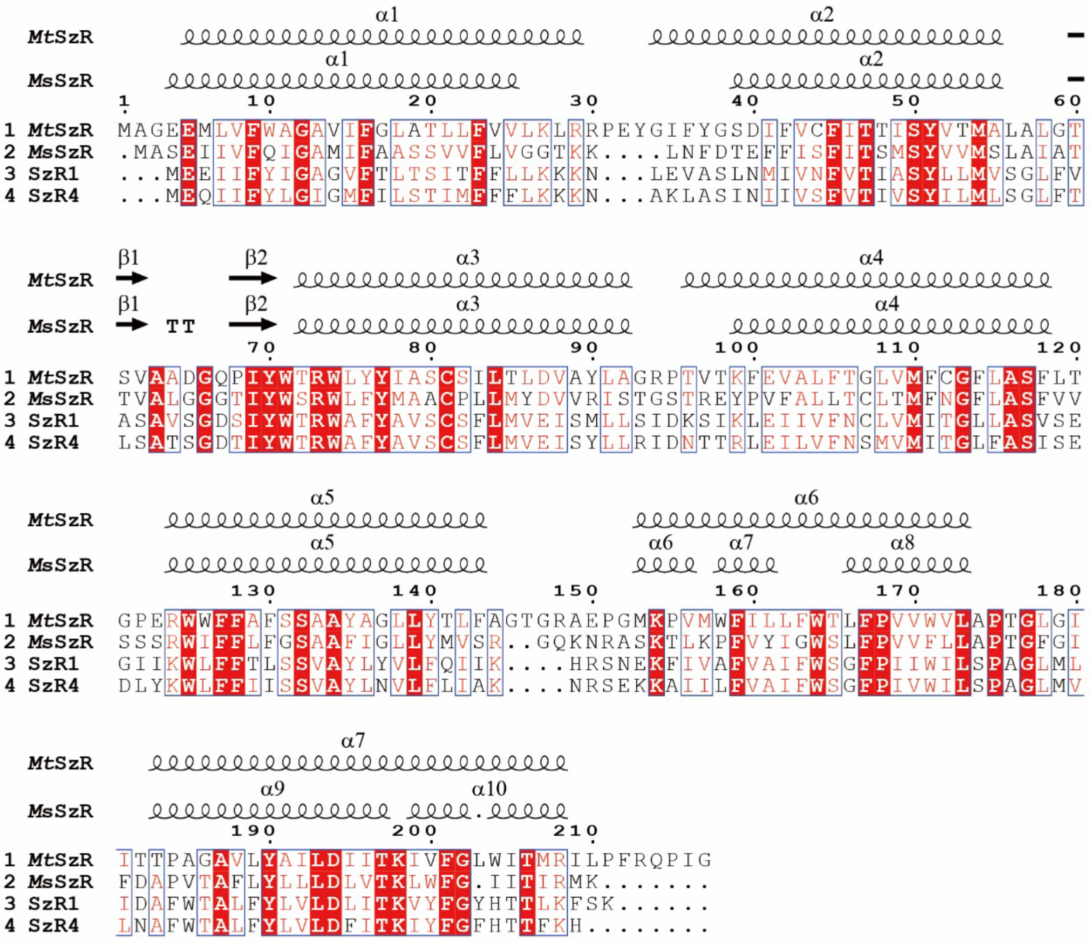


**Figure S4.** Alignment of amino acid sequences of SzRs.

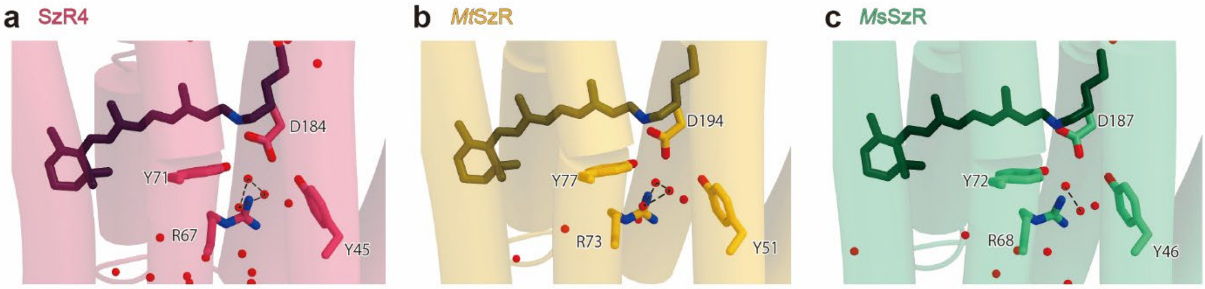


**Figure S5.** Comparison of RSB. Black dashed lines indicate hydrogen bonds. **a**. SzR4. **b**. *Mt*SzR. **c**. *Ms*SzR.

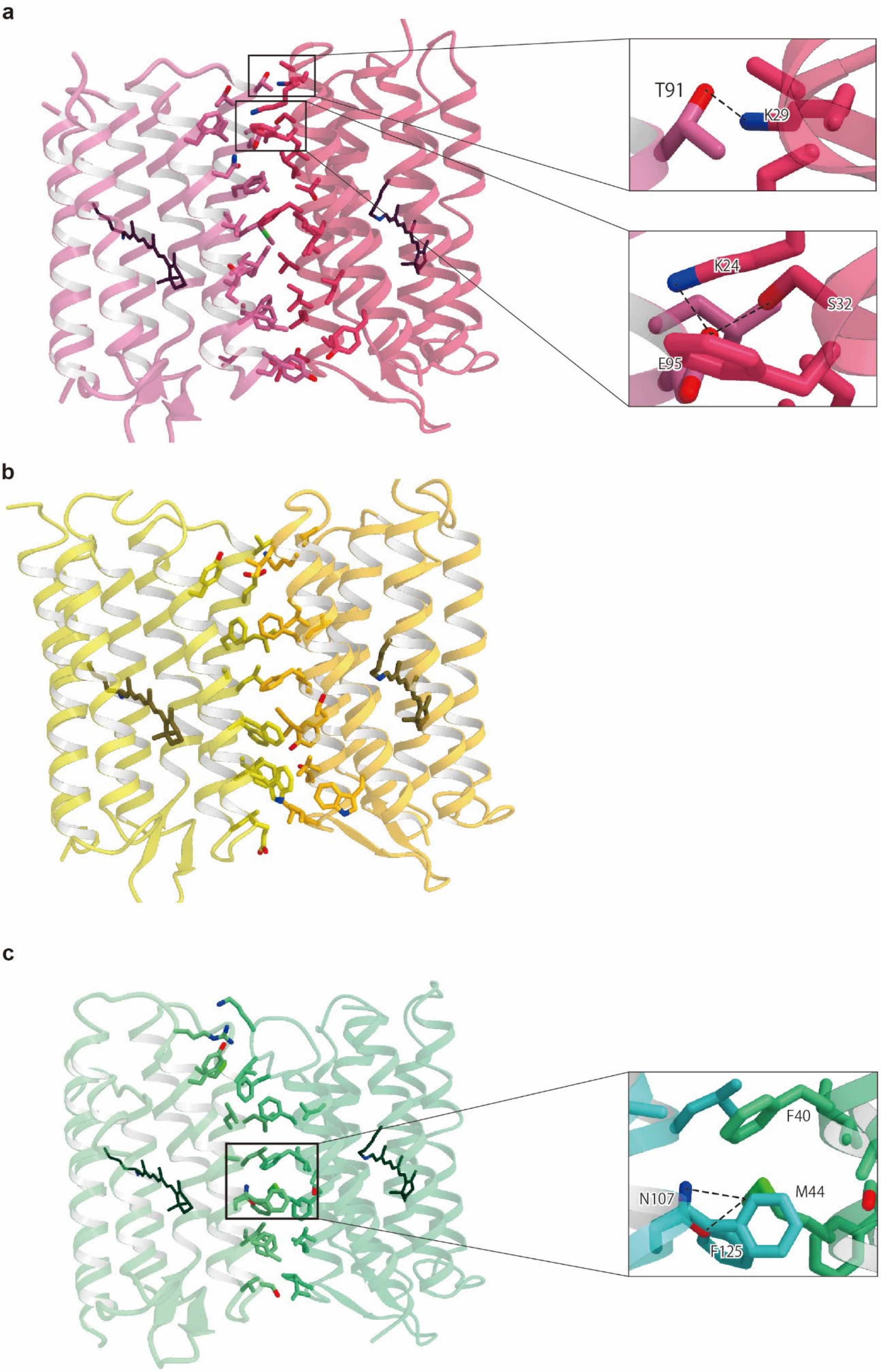


**Figure S6.** Comparison of trimer interface. Black dashed lines indicate hydrogen bonds. **a**. SzR4. **b**. *Mt*SzR. **c**. *Ms*SzR.

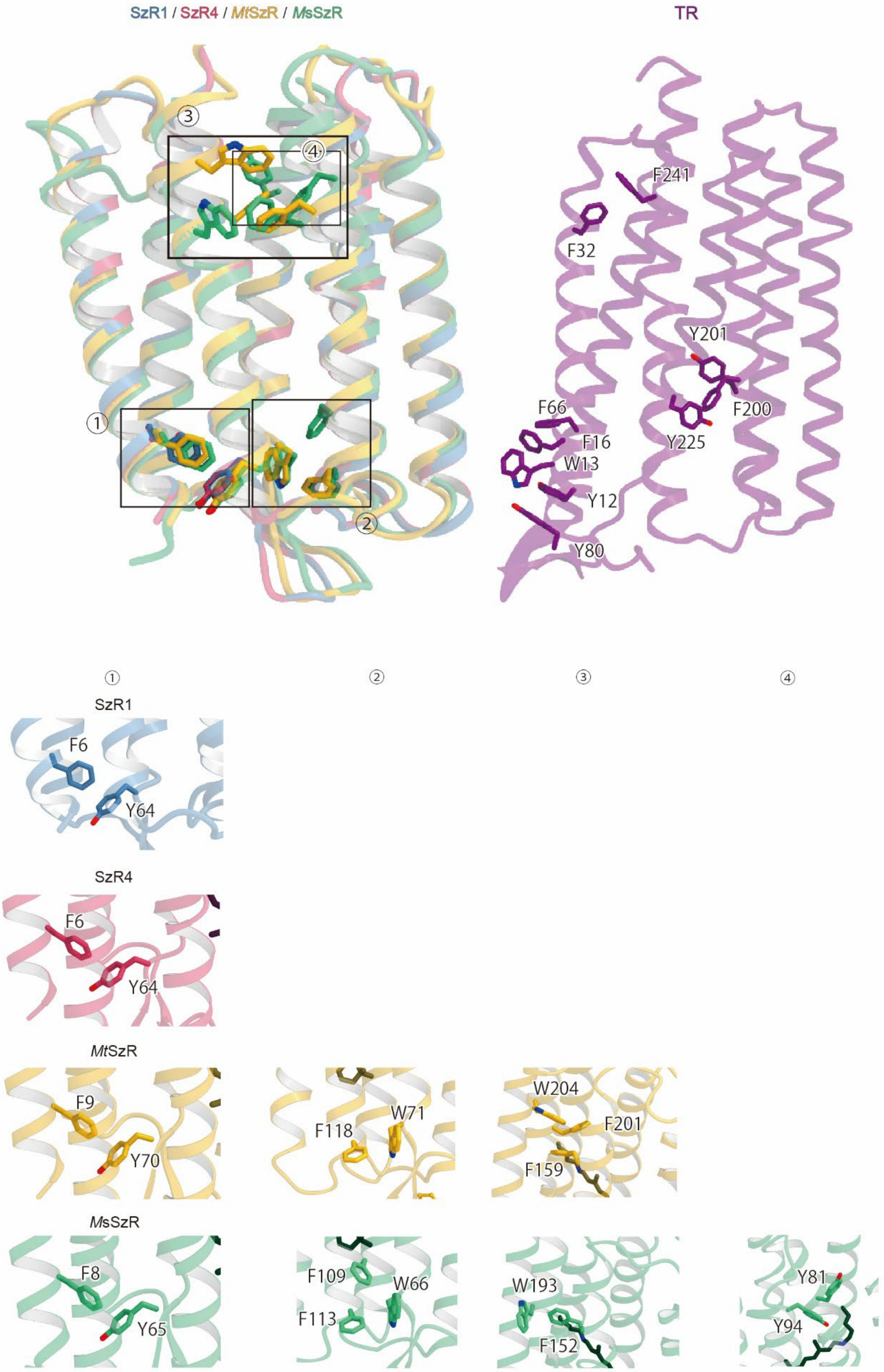


**Figure S7.** Comparison of the helix-to-helix aromatic interactions of SzRs. Compared to SzR4, both *Mt*SzR and *Ms*SzR exhibit a greater number of interhelical interactions involving aromatic residues.

| | *Mt*SzR | *Ms*SzR |
|---|---|---|
| PDB | XXXX | XXXX |
| EMDB | EMD-XXXXX | EMD-XXXXX |
| **Data collection** | | |
| Microscope | Titan Krios (Thermo Fisher Scientific) | |
| Voltage (kV) | 300 | |
| Detector | Gatan K3 camera (Gatan) | |
| Magnification | ×105,000 | |
| Electron dose (e−/Å) | 50 | |
| Defocus (μm) | −0.6 ~ −1.6 | |
| Pixel size (Å/pix) | 0.83 | |
| Number of movies | 12,309 | 9,891 |
| Picked particles | 8,692,253 | 7,273,694 |
| Final particles | 145,730 | 187,471 |
| Map resolution (Å) | 2.38 | 2.67 |
| FSC threshold | 0.143 | |
| **Model refinement** | | |
| Atoms | 5943 | 5100 |
| R.M.S.D. from ideal | | |
| Bond length | 0.0086 | 0.0040 |
| Bond angle | 0.72 | 0.52 |
| Validation | | |
| Clashscore | 2.47 | 5.24 |
| Rotamer outliers (%) | 0.00 | 1.96 |
| Ramachandran plot | | |
| Favored (%) | 99.04 | 99.17 |
| Allowed (%) | 0.96 | 0.83 |
| Outlier (%) | 0 | 0 |

**Table S1.** The conditions for single particle analyses using cryo-electron microscopy and the statistics of the obtained maps and models are shown. Structural analyses were performed using cryoSPARC, and modeling was performed using COOT, PHENIX. Models were evaluated using MolProbity.
